# On the accuracy of methods identifying gait events using optical motion capture and a single inertial measurement unit on the sacrum

**DOI:** 10.1101/2025.03.09.642234

**Authors:** Vinicius Cavassano Zampier, Morten Bilde Simonsen, Fabio Augusto Barbieri, Anderson Souza Oliveira

## Abstract

Gait events (i.e., heel strikes and toe-offs) are essential for extracting spatiotemporal parameters and segmenting biological signals. While force platforms and optical motion capture (OMC) are ideal for identifying such events, inertial measurement units (IMUs) are cheaper and more applicable, especially for applications outside traditional lab settings. This study aimed to compare IMU- and OMC-based gait event detection to force plates. Seventeen adults walked on a walkway, stepping on two force plates while an IMU on the sacrum and retro-reflective markers on the calcaneus and 5th metatarsus captured foot kinematics. Gait events were identified using two OMC and two IMU methods (OMC1, OMC2, IMU1, IMU2). OMC1 detects gait events using vertical marker velocity shifts, OMC2 uses sagittal velocity thresholds, IMU1 applies wavelet-based differentiation and IMU2 identifies peaks in acceleration signals. Heel strikes and toe-offs were compared to force plate data, assessing root-mean-square error (RMSE), and intra-subject coefficient of variation (CoV). For heel strike events, OMC1 presented the lowest RMSE (∼14.3ms), significantly differing from IMU1 (RMSE: 50.6ms; p<0,001) and IMU2 (RMSE: 61.1ms; p<0,001). For toe-offs, OMC1 presented the lowest RMSE (∼17.3ms), differing from IMU1 (54.7ms; p<0,001) and IMU2 (74.8ms; p<0,001). IMU2 also showed the highest CoV (∼23.9ms) differing from OMC1 (∼7.1ms; p<0,001) and IMU1 (9.2ms; p<0,001). Lower accuracy and greater variability in IMU methods may stem from the approach used to detect gait events. Thus, OMC methods more accurately detect gait events than sacrum-mounted IMUs. While IMUs offer an alternative, researchers should be cautious of their accuracy and variability limitations.

## Introduction

The identification of gait events such as heel strikes and toe-offs during walking allows the extraction of clinically relevant spatiotemporal parameters (Gaßner et al., 2020; Tabard-Fougere et al., 2022; Ditunno J., & Scivoletto G., 2009). Step length and cadence allow for assessing walking rhythm and potential asymmetries that are markers of neurodegenerative diseases such as Parkinson’s disease (Faria et al., 2023; Godi et al., 2021). Double support periods illustrate balance control, as prolonged double support may indicate increased instability and/or fall risk (Vitorio et al., 2023). Therefore, accurately identifying gait events is crucial to evaluating motor function and the overall status in clinical populations.

The gold-standard method of defining gait events requires force plate measurements of ground reaction forces (GRF). The start and end instants of touching the floor are typically detectable through the change in vertical GRF (vGRF) from a baseline unloaded measurement when a person steps onto the plate (Marasovic et al., 2009; Samson et al., 2001). Another laboratory-based method to detect gait events involves motion capture using markers placed over the calcaneus and metatarsal foot bones to track vertical and forward foot positions (Ghoussayni et al., 2004). The accuracy of markers in identifying gait events has been shown to vary between 30 and 100 milliseconds depending on the position of the markers (Ghoussayni et al., 2004; Zeni et al., 2008; Caron-Laramée et al., 2023; Zahradka et al., 2020). However, optical motion capture is a technique limited to laboratory settings.

Plantar pressure insoles offer an alternative for detecting gait events outside laboratories, identifying foot pressure similarly to vGRF (Salis et al., 2021). However, high cost limits their practicality for clinical use. Inertial measurement units (IMUs) containing accelerometers, gyroscopes, and magnetometers are a cost-effective, lightweight, and easy-to-apply tool for gait analysis (Godfrey et al., 2014; Del Din et al., 2015). Recently, gait event detection has been carried out using IMU data mimicking smartphone usage (Larsen et al., 2024), showing its potential for remote monitoring. While IMUs are a promising method for gait analysis in clinical studies, the accuracy of their gait event detection remains unclear. Previous research has used a single sacrum-mounted IMU to measure step length and walking speed (Del Din et al., 2016;

De-Oliveira et al., 2020), showing strong correlations (0.84<r<0.99) between predicted and experimental variables extracted from motion capture systems and pressure sensors carpet. Other IMU-based studies investigated gait using multiple accelerometers placed on the sacrum and ankles, reporting correlation between 0.82 to 0.99 when compared with OMC systems (Hundza et al., 2014; Kluge et al., 2017). Moreover, a sacrum-mounted IMU-based method has the advantage of identifying bilateral gait events by utilizing the different motion planes, highlighting its applicability in studies on gait asymmetry (Zijlstra, & Hof., 2003; McCamley et al., 2012). However, these validation procedures extract the final gait parameters without determining the accuracy of the heel strike/toe-off detection, while also using surrogates of the gold standard to assess ground truth, such as pressure mats (Del Din et al., 2015).

Since IMUs placed on the sacrum are an alternative to detect gait events, it is possible to use such events in further analysis involving the segmentation of surface electromyography (EMG) and/or electroencephalography (EEG) recorded during walking. However, inconsistent/incorrect gait event detection can substantially compromise EMG/EEG data analysis. For instance, a consistent delay in detecting initial contact can be incorporated into the bio-signal, shifting events in time and leading researchers and readers to misinterpretations. Therefore, this study aimed to determine the quality of gait event detection using single IMUs compared to optical motion capture (OMC) against the gold-standard force plate measurements.

## Methods

### Participants

Seventeen participants (16 male/1 female, age: 31±8 years, height: 179±6 cm, body mass: 78±7 kg) without previous neurological or musculoskeletal injuries preventing treadmill walking participated in the present study. Participants provided verbal and written informed consent. All methods followed the Declaration of Helsinki (2004) and were approved by the local Research Ethics Committee (Case number: 2023-505-00139).

### Experimental design and instrumentation

Participants were instructed to walk at their preferred speed on an 8-m-long walkway in a single session. At the center of the walkway were two force plates recording tri-dimensional GRF (BMS400600, AMTI, Watertown, USA), 1,000 Hz sampling frequency). Participants were instructed to step onto each force plate with one foot, with the option to use either foot for the first and second plates. They walked at a preferred speed for 60 seconds along the walkway, stepping onto the force plates in both directions while wearing their own footwear. A total of five to seven 60-second trials were recorded for each participant.

Before data recordings, participants were fitted with tri-dimensional IMUs (Trigno Avanti Sensor, Delsys Inc., Massachusetts, USA) that recorded acceleration at ±16g and a 1,259 Hz. The sensors were placed on the participant’s sacrum following recommendations from the literature (Godfrey et al., 2014; Del Din et al., 2016). In addition, we acquired the tri-dimensional position of retro-reflexive markers using an OMC (12-camera QTM 2023.2, Qualisys, Göteborg, Sweden, 100 Hz sampling frequency). The markers were placed over the participant’s walking shoes to represent their calcaneus and first metatarsal bilaterally. The GRF and IMU data were recorded/synchronized through the motion capture software (QTM 2023.2, Qualisys, Göteborg, Sweden).

### Data processing

The vGRF was used as the gold-standard method to identify gait events. The timing of the heel strike and toe-off events were defined using the raw vGRF, using a force threshold of 15 N for both force plates. The calcaneus and metatarsal marker data were interpolated to match the 1000 Hz sampling frequency and subsequently low-pass filtered using a 4^th^-order Butterworth (15 Hz cut-off frequency). We used two methods to identify gait events using OMC. In the first (OMC1, Zeni et al 2008), The heel strike events were determined as the point at the calcaneus marker vertical signal shifts from a positive to a negative vertical direction. The toe-off events were determined when the vertical velocity of the 1st metatarsal marker shifted from negative to positive vertical direction. In the second method (OMC2), we used the method described in Ghoussayni et al., 2004. We first low-pass filtered the marker’s data (4^th^ order Butterworth filter, 10 Hz cut-off frequency). Next, markers’ velocities in the sagittal plane were calculated to determine the heel strike and toe-off events when the marker stopped moving in vertical and antero-posterior directions. Subsequently, the average and standard deviation of the markers’ sagittal velocities during the contact periods were calculated using a threshold value to account for low-amplitude marker movements. We adapted the proposed threshold from 50 ms to 75 ms for detecting toe-offs, as it provided better results using the marker on the first metatarsal.

Accelerometry data were down-sampled to 1,000 Hz to match the vGRF data. Subsequently, the data was low-pass filtered using a 4^th^-order Butterworth (30 Hz cut-off frequency). We used two methods to define gait events. The first (IMU1, Deldin et al. 2016), followed five steps: i) numerical integration of sacrum vertical acceleration; ii) integrated signal differentiation using the continuous wavelet transformation, resulting in the first differentiated signal, this is a decomposition technique which allows the temporal localization of cyclical events in non-stationary signals (McCamley et al., 2012); iii) define heel strike by finding the local minima times of the first differentiated signal; iv) differentiation of first differentiated signal using continuous wavelet transformation resulting in second differentiated signal; v) define toe-off by finding the local maxima times of the second differentiated signal. The authors advise a correction of the vertical acceleration to horizontal-vertical frame. However, this correction did not yield superior results compared to the non-corrected data and was not used in our results. In the second method (IMU2, Weersink et al. 2021) we followed three steps: i) remove any gravity or impulse-like artifacts and define the time points of maximum positive peaks of vertical acceleration signal consider a window 0.35 seconds between each peak; ii) identify the toe-offs by finding the first negative peak within a window of 0.15 seconds after the maximum positive peak defined on the first step; iii) defining the heel strikes by finding the first negative peak 0.15 seconds before the largest positive peak in the antero-posterior acceleration signal.

An average of 92±17 stance periods per participant were analyzed, including force plate data and corresponding stance periods from OMC1, OMC2, IMU1, and IMU2 methods for both the right and left legs, totaling 1598 steps. A total of 92±17 stance periods with force plate and respective stance periods from OMC1, OMC2, IMU1 and IMU2 methods per participant across both right and left leg were analyzed (1598 steps in total). Since there were no relevant differences between body sides on a preliminary analysis, results from both legs were merged. We computed the root-mean-square error (RMSE) and mean absolute error (MAE) by comparing the gold-standard force-platform gait event detection with OMC1, OMC2, IMU1 and IMU2. Moreover, we computed the intra-subject coefficient of variation (CoV) to illustrate intra-subject variability in the gait event estimations. CoV was defined as:

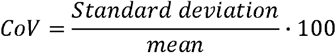

Finally, the stance phase time was computed for all methods as the time difference between a heel strike and the preceding toe-off for the same side.

### Statistical analysis

Statistical analyses were performed using SPSS 20.0 (SPSS, Inc.) software, and the significance level was set to α = 0.05. Normality and sphericity were verified through Shapiro-Wilk and Mauchly’s test on each dependent variable (stance time, stance time CoV, stance time RMSE, and range of prediction errors). When necessary, sphericity was corrected using Greenhouse-Geisser procedures. To assess the effect of gait detection methods (force plate, OMC1, OMC2, IMU1, and IMU2) on stance time and stance time CoV. Repeated measures one-way ANOVA with Bonferroni post-hoc corrections when applicable. We also used repeated measures one-way-ANOVA to verify the accuracy across methods regarding stance RMSE and range of prediction errors with the Bonferroni post-hoc correction to explore the differences between the force plate and the other methods (OMC1, OMC2, IMU1, and IMU2). The effect sizes for the ANOVA results were interpreted using partial eta-squared (η^2^), with thresholds of 0.01, 0.06, and 0.14 indicating small, medium, and large effects, respectively (Richardson., 2011). Pearson correlations were performed to verify the correlation between OMCs and IMUs stance times with the force plate measurement. The strength of the correlation was classified based on the absolute value of Pearson’s r as follows: low (r≤ 0.3), moderate (0.4< r ≤ 0.6), and high (r > 0.6) (Akoglu., 2018). Bland-Altman plots were used to compare the timing of gait events obtained using the implemented algorithms against the ground truth values from force plates. Accuracy was defined as the mean difference between two methods of measurement, and 95% limits of agreement were calculated to assess the variability of the differences.

## Results

We reviewed the identified gait events and excluded heel strike/toe-off pairs with an absolute error greater than 250 ms, likely due to erroneous steps during data collection. Therefore, we used 90±19 steps from each participant. The computed error for heel strikes and toe-offs calculated using OMC and IMU methods from each participant are shown in Figure 1. Generally, gait events extracted from OMC methods (left panels) presented errors within ±50 ms. Both IMU methods presented larger errors and greater intra-subject dispersion across the identified events. Notably, heel strike events were typically identified earlier than the actual event, while toe-offs were identified later than the true events, especially for the IMU methods.

**Figure 1.**
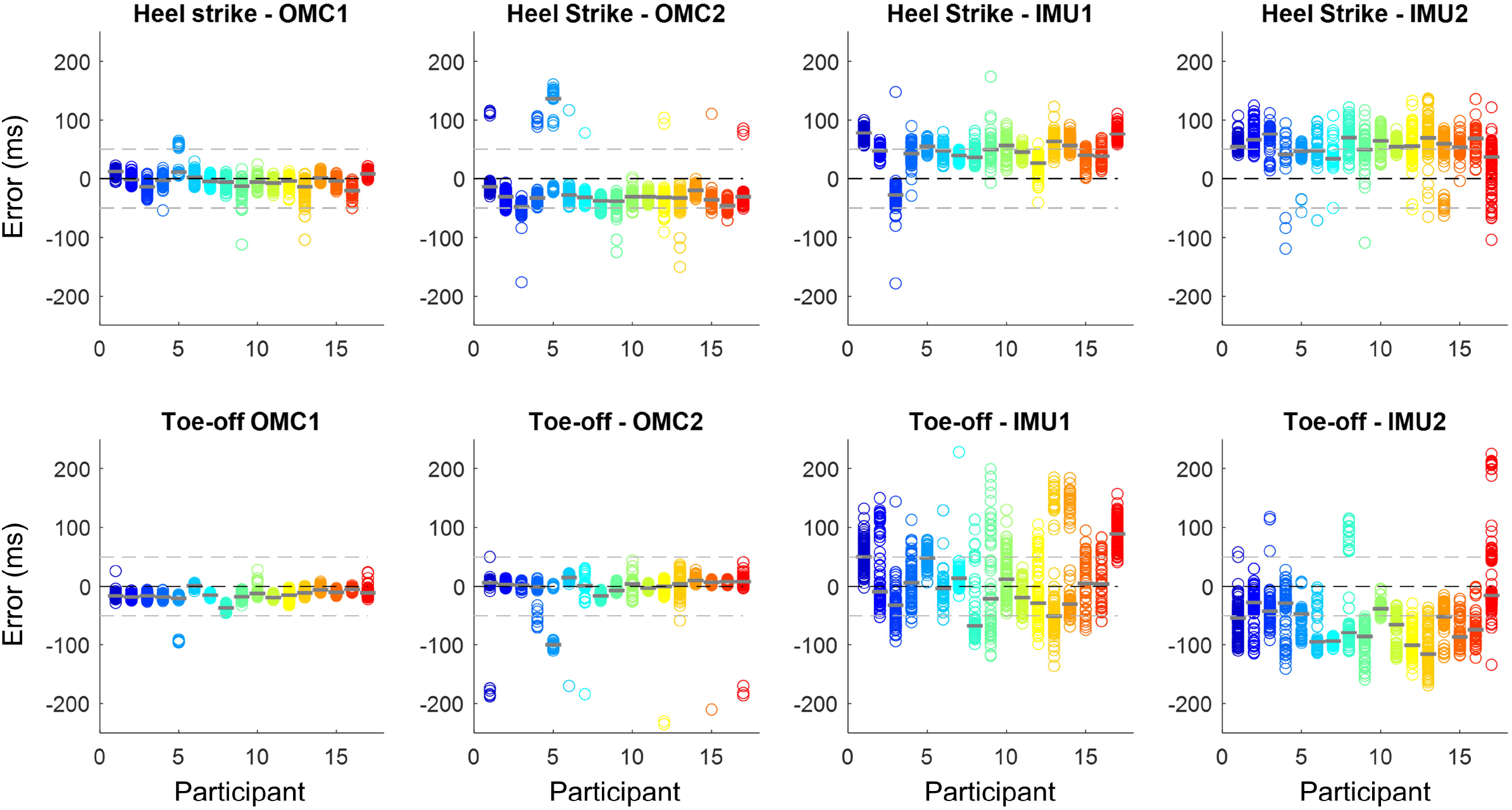
Absolute error (in ms) in heel strike and toe-off identification using two OMC (OMC1 and OMC2) and two IMU methods (IMU1 and IMU2) when compared to the gold-standard identification using force plates. Each circle represents a gait event, and each color represents a participant (N = 17), with the median error represented by the gray line over the circles. Horizontal dashed lines represent no error (0 ms), 50 and -50 ms.

### Prediction quality from OMC and IMU

There was a significant main effect of methods for RMSE for both heel strikes (F_3,48_=39.079; p<0.001; ηp^2^=0.71) and toe-offs (F_3,48_=39.658; p<0.001; ηp^2^=0.71). OMC1 (14.3±9.0 ms) to identify heel strikes presented lower error when compared to OMC2 (42.1±19.2 ms, p<0.001), IMU1 (50.6±13.9 ms, p<0.001) and IMU2 (61.1±10.1 ms, p<0.001, Figure 2A). To identify toe-offs, the OMC1 (17.3±19.3 ms) presented lower error when compared to the IMU1 (54.7±21.4 ms, p<0.001) and IMU2 (74.8±20.6 ms, p<0.001). The OMC2 (21.6±19.37 ms) presented lower error when compared to the IMU1 (p=0.002) and IMU2 (p<0.001, Figure 2C).

**Figure 2.**
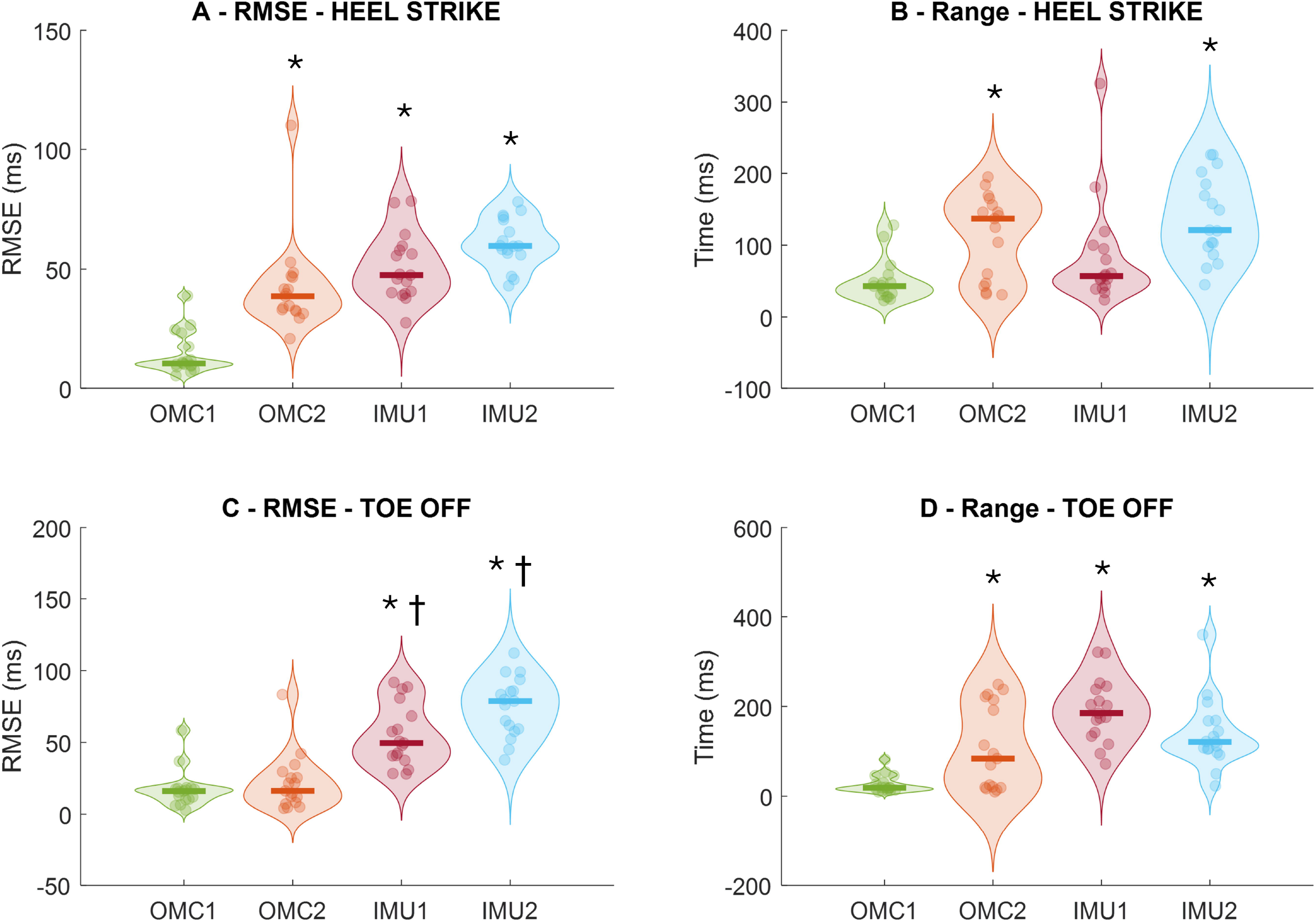
Root mean square error (RMSE) of heel strike (A) and toe-off (C), and intra-subject prediction range if heel strike (B) and toe-off (D) from the comparison of force plate and methods from optical motion capture (OMC) and inertial measurement units (IMUs) to detect heel strikes (upper row) and toe-off events (bottom row). * denotes significant difference in relation to OMC1 (p<0.01); † denotes significant different in relation to OMC2 (p<0.05).

Regarding the range of prediction errors, there was a main effect of method for both heel strikes (F_3,48_=10.131; p<0.001; ηp^2^=0.39) and toe-offs (F_3,48_=15.224; p<0.001; ηp^2^=0.48). OMC1 (49.7±29.4 ms) presented lower range of prediction errors to identify heel strikes when compared to OMC2 (112.6±58.5 ms, p=0.004) and to IMU2 (138.2±58.0 ms, p<0.001, Figure 2B). Regarding toe-offs, OMC1 (27.0±19.3 ms) presented lower range of prediction errors when compared to OMC2 (107.7±94.0 ms, p=0.013), IMU1 (191.5±70.0 ms, p<0.001), and IMU2 (138.0±76.3 ms, p<0.001, Figure 2D).

The Bland-Altman plots (Figure 3) demonstrated a smaller range on the limits of agreement for OMC1 (0.14 s, Figure 3A) and OMC2 (0.12 s, Figure 3B) when compared to the IMU1 (0.28 s, Figure 3C) and IMU2 (0.73 s, Figure 3D). Also, participants’ stance time estimated in IMU2 follow a trend with a slope, suggesting a potential systematic bias related to amplitude indicating a possible overestimation or underestimation of certain values rather than random variation.

**Figure 3.**
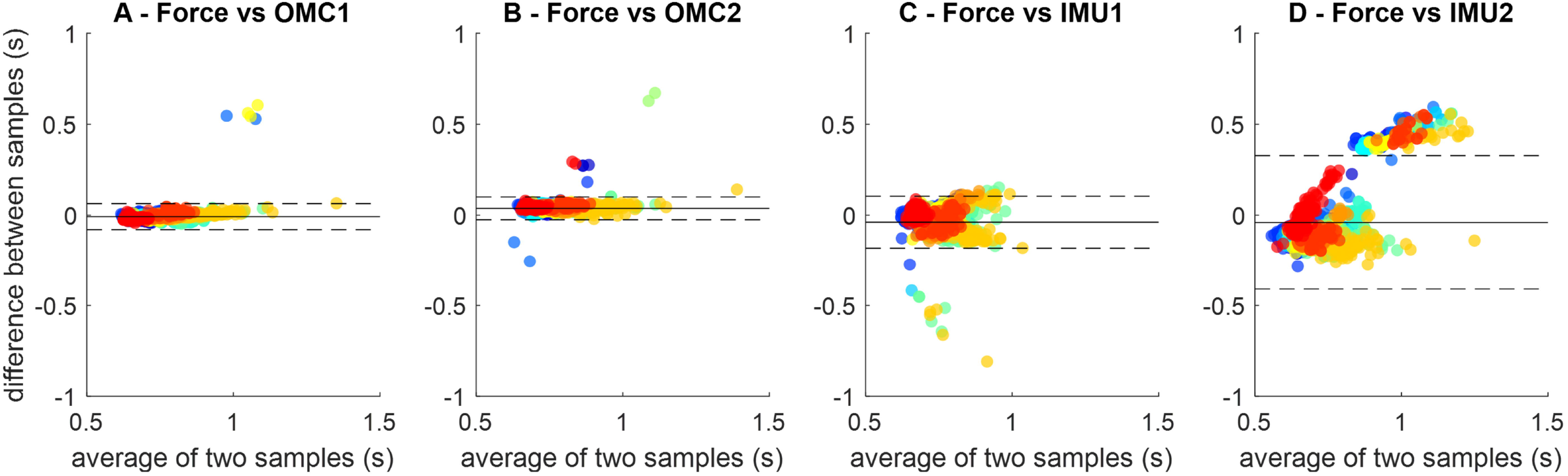
Bland-Altman plots of all stance periods were identified across all participants (identified as different colors).

### Stance time across methods

There was a main effect of method for stance time (F_4,64_=12.055; p<0.001; ηp^2^=0.43, Figure 4A). The stance time extracted from the force plates (0.76±0.01 s) was lower than OMC2 (0.85±0.02 s, p=0.045) and higher than IMU1 (0.73±0.01 s, p=0.045). The stance extracted from OMC2 was higher from IMU1 (p=0.001) and IMU2 (0.739±0.025s, p=0.018). The stance phase time extracted from the OMC1 was overestimated by only 0.26±5.5%, whereas OMC2 was overestimated by 11.4±15.3% (Figure 4A). IMU1 provided stance phases underestimated by 4.7±3.8%, whereas IMU2 presented underestimations at 3.9±8.5%.

**Figure 4.**
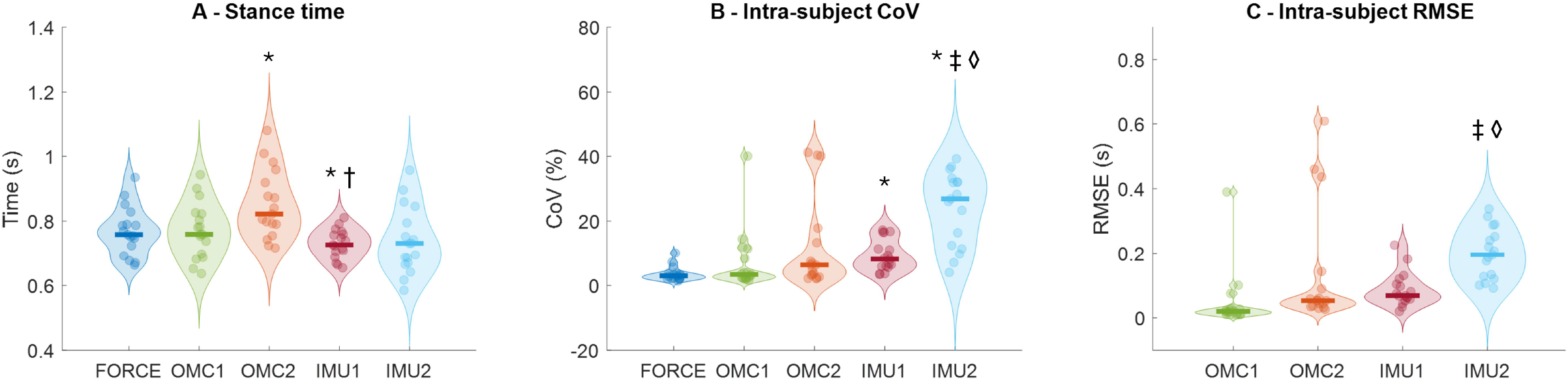
Stance time (A), stance time intra-subject coefficient of variation (CoV, B) and stance time intra-subject root mean square error (RMSE, C) extracted from force plate data and different methods using optical motion capture (OMC) and inertial measurement units (IMU). * denotes significant difference in relation to Force (p<0.01); † denotes significant different in relation to OMC2 (p<0.05); ‡ denotes significant different in relation to OMC1 (p<0.05). * denotes significant difference in relation to OMC’ (p<0.05).

### Intra-subject coefficient of variation (CoV)

There was a main effect of method on CoV (F_4,64_=12.919; p<0.001; ηp^2^=0.44, Figure 4B). The CoV was higher in IMU1 (9.2±1.2, p<0.001) and IMU2 (23.9±2.7, p<0.001) when compared to the force plate (3.5±0.5). Moreover, the IMU2 presented a higher CoV when compared to OMC1 (7.1±2.2, p<0.001) and IMU1(p=0.001). The CoV from the force measurement was the lowest on average, below 4% (3.5±2.2%). Both OMC methods presented average variability between 7% and 12%, while IMU methods presented averages between 9% and 24%, with greater inter-subject fluctuations, especially for IMU2.

### Intra-subject root mean square error

There was a main effect of method for RMSE (F_3,48_=6.105; p=0.010; ηp^2^=0.27, Figure 4C). The RMSE extracted from OMC1 (0.048±0.02 s) was significantly lower when compared to IMU2 (0.199±0.019 s, p<0.001). Moreover, the RMSE from IMU1 (0.087±0.013 s) was significantly lower when compared to IMU2 (p<0,001). The RMSE calculated within a single participant’s data demonstrated that OMC methods yield errors predominantly equal to or below 0.1 s, considering that both methods presented outliers. Regarding IMU methods, their errors were, on average around 0.15 s.

### Correlation between force and OMC/IMU stance phase times

The stance times extracted from the force measurements were highly associated with those extracted from the OMC1 and IMU1 (Figure 5A and 5C, r>0.85, p<0.001). In contrast, the association was slightly weaker for IMU2 (r = 0.76, p<0.001, Figure 5D), and OMC2 association presented an outlier that reduced the Pearson’s correlation to 0.37 (p = 0.13, Figure 5B). Removing the outlier participant would increase the association to a significant 0.74 (p<0.001).

**Figure 5.**
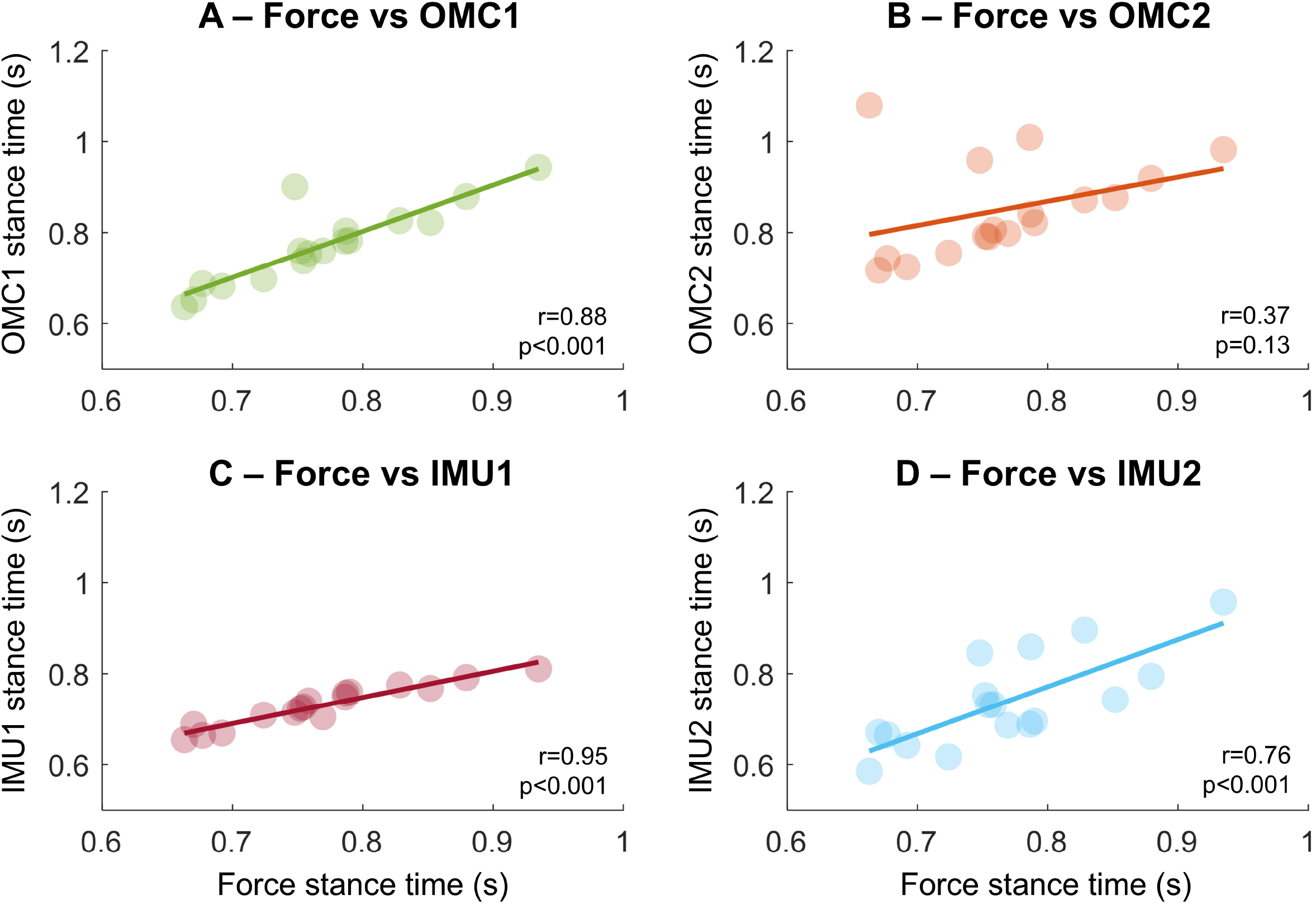
Association of the stance phase time extracted using gold-standard force measurements compared with two methods based on optical motion capture (OMC, A and B) and two methods using IMU sensors placed on the sacrum (in C and D).

## Discussion

The main findings of the present study were that gait events identified using OMC methods reach average errors as low as 15 ms, whereas the best IMU detection reaches ∼50 ms errors. However, IMU detections present greater intra-subject variability, especially for toe-off events. Calculations of stance time revealed that the RMSE from OMC1 (48±9 ms) was substantially lower when compared to IMU1 (87±5 ms) and especially IMU2 (199±7 ms). Moreover, stance time estimations from IMUs generate greater stride-to-stride variability than force and OMC methods. Nonetheless, strong associations were found between stance times extracted from the force and both OMC and IMU methods. These results demonstrate that OMC generates gait events and stance phase estimations with lower errors and lower variability, and researchers interested in using sacrum-mounted IMUs for gait analysis must be highly cautious with their analysis/interpretation due to larger errors and inherent variability.

### Prediction errors from OMC methods

Previous studies have reported absolute errors in predicting gait events from OMC methods ranging between 4-30 ms, which have been deemed acceptable (Caron-Laramée et al., 2023; Zahradka et al., 2020; Harsen et al., 2002). Zeni et al. (2008) reported average errors between 20-25 ms in identifying gait events in young adults following the published OMC-based methods (Ghoussayni et al., 2004; Zahradka et al., 2020). Caron-Laramée et al. (2023) observed similar values, which compared an OMC-based method with a force plate. In pathological populations such as Parkinson’s disease, gait event errors were 30 ms compared to events extracted from young adults using OMC (Bonci et al., 2022). The investigated OMC-based algorithms presented an average absolute error of ∼25 ms, reinforcing the quality and robustness of OMC-based gait detection methods (Zeni et al., 2008). Also, we observed that OMC methods presented lower intra-subject variability (∼7% to 12%) than IMUs (∼9% to 12%), reinforcing the greater reliability of the OMC method, as it indicates that detection errors remain more consistent.

### Prediction errors from IMU methods

Several studies have been using different IMU methods to detect gait events. Boutaayamou et al. (2015) achieved an error of approximately 15 ms compared to markers data in detecting gait events using two accelerometers placed on the calcaneus and the fifth metatarsal. While Brahimetaj et al. (2024) used a single sacrum-mounted IMU, they observed that the error can reach around 40 ms compared to other IMU methods. Studies placing IMUs on participants’ shanks (Romijnders et al., 2021) or the foot dorso (Voisard et al., 2024) have reported drastically reduced errors (∼8-14 ms) when compared to OMC methods, demonstrating that multi-sensor approaches for gait event detection are more accurate. However, more sensors will reduce the potential for real-world applications of the technique, due to the need to fix multiple sensors. Moreover, methods based on data from IMUs fixed on the shanks and/or feet require bilateral fixation of sensors, while the sacrum-mounted methods require only one accelerometer (McCamley et al., 2012).

The lower accuracy and high variability in detecting gait events from IMU-based methods urge caution when applying such methods for bio-signal segmentation (e.g., EEG or EMG), which might be temporally misaligned and contain random variability across gait cycles. The lower prediction quality from sacrum-mounted IMU methods may be related to the temporal mismatch between the actual gait events and the available traceable events within the IMU data. Peaks/valleys from IMU data may indicate when the foot touches the floor, but the two events are not fully temporally aligned since there is a chain of mechanical events from the true foot touch to the response in the IMU data. Moreover, some IMU methods perform signal transformations (e.g., wavelets) to improve such event detection, but the transformed data might still be temporally misaligned with the real events. Finally, tri-dimensional accelerometry/gyroscopic data is highly variable across participants, whereas foot marker trajectories during walking present substantially reduced inter-subject variability. Regardless of the limitations of IMU methods, they did not influence inter-subject patterns in stance time estimation when stance times are averaged, as demonstrated by the strong correlation between experimental and predicted data. Godfrey et al. (2015) observed a high intraclass correlation coefficient (0.84) between stance times extracted from a sacrum-mounted IMU method and pressure mat. This might be attributed to the estimated parameters data averaged across ∼60 cycles defined by steps. Likewise, our study averaged ∼60 overground steps/participant, substantially improving the robustness of the average.

## Conclusion

In conclusion, OMC and IMU methods offer complementary advantages depending on the context of the application. IMUs, despite their lower accuracy, provide a practical and cost-effective solution for real-world gait monitoring if their limitations are acknowledged and carefully considered during data interpretation. Our results demonstrated that OMC1 had the highest accuracy in detecting gait events compared to IMU1 and IMU2. IMU2 presented the highest variability, systematic bias, and stance time estimation errors, whereas OMC1 and IMU1 presented strong correlations with force plate data. This study reinforces the importance of rigorous validation and calibration of gait events tracking methods to detect heel strikes and toe-offs reliably. Future research should focus on developing gait event detection algorithms that apply machine learning methods to improve accuracy and reduce variability, possibly expanding the user groups to diverse clinical populations.

## CRediT authorship contribution statement

### Vinicius Cavassano Zampier

Writing – review & editing, Writing – original draft, Methodology, Data curation, Conceptualization, Formal analysis. **Morten Bilde Simonsen**: Writing – review & editing, Data curation, Methodology. **Fabio Augusto Barbieri**: Writing – review & editing, Funding acquisition, Supervision. **Anderson Souza Oliveira**: Writing – review & editing, Writing – original draft, Methodology, formal analysis, Resources, Conceptualization, Supervision, Project administration.

## Acknowledgments

We would like to thank all the participants who volunteered for this study.

